# Low-Intensity Ultrasound Stimulates TAZ in Schwann cells

**DOI:** 10.1101/2025.05.08.652925

**Authors:** Jenica Acheta, Haley Jeanette, Amanda S. Mondschein, Meghana Lanka, Sophie Belin, Yannick Poitelon

## Abstract

Mechanosensation, the ability of cells to detect and respond to mechanical forces by transducing them into biochemical signals, is essential for various cellular processes, including morphogenesis, development, tissue homeostasis, and response to injury. In the peripheral nervous system (PNS), Schwann cells play a critical role in nerve development, myelination, and regeneration. These cells are highly responsive to mechanical cues such as tension, compression, and shear forces, which influence their fate, proliferation, differentiation, and regenerative capacity. In this study, we demonstrate that *in vitro* application of Low Intensity Ultrasound (LIU) transiently increases Schwann cell proliferation. Notably, our results show that LIU selectively activates TAZ, but not YAP, both nuclear transducers of the Hippo pathway. Additionally, we show that the LIU treatment upregulates nerve growth factor (NGF) expression in both Schwann cells and sensory neurons, suggesting a role for LIU in promoting neurotrophic support. This study highlights LIU as a mechanotherapeutic tool that enhances intrinsic regenerative functions in Schwann cells, such as neurotrophic support to neurons via NGF.

**Main Points:**

- Application of LIU promotes SC proliferation and production of nerve growth factors.
- TAZ is activated in Schwann cells following LIU application.

## 1. Introduction

Mechanosensation, the ability of cells to detect and respond to mechanical forces, plays a fundamental role in development, tissue homeostasis, and injury responses (Di et al. 2023). In the peripheral nervous system (PNS), Schwann cells play a fundamental role in nerve development, myelination, and regeneration. Given their close association with axons and the extracellular matrix (ECM), Schwann cells are highly sensitive to mechanical cues, including tension, compression, and shear forces (Belin, Zuloaga, and Poitelon 2017). These forces influence crucial aspects of Schwann cell function, including their proliferation, differentiation, myelination, and regenerative capacity after nerve injury (Belin, Zuloaga, and Poitelon 2017). Understanding how Schwann cells integrate mechanical signals with biochemical pathways has become an area of growing interest, as it has significant implications for peripheral neuropathies, nerve repair strategies, and myelin-related disorders.

Schwann cells rely on a variety of mechanotransduction mechanisms to sense and respond to their mechanical environment. One of the primary mechanisms involves mechanosensitive ion channels such as Piezo channels, which detect mechanical forces and regulate downstream cellular processes. Activation of Piezo channels in Schwann cells leads to calcium influx, influencing their proliferation, differentiation, and myelination (Acheta, Bhatia, et al. 2022; Suttinont et al. 2024). Additionally, Schwann cells interact extensively with the ECM via integrins (Feltri et al. 2021), which act as mechanosensitive receptors that relay external mechanical signals into intracellular responses in part through striatins (Weaver et al. 2025). These integrin-ECM interactions regulate focal adhesion dynamics, cytoskeletal remodeling, and downstream signaling pathways critical for Schwann cell migration, radial sorting, and myelination (Berti et al. 2006).

Among the key intracellular pathways influenced by mechanical signals, the Hippo-YAP/TAZ signaling axis has emerged as a central regulator of mechanotransduction in Schwann cells. YAP (Yes-associated protein) and TAZ (transcriptional coactivator with PDZ-binding motif) are mechanosensitive transcriptional regulators that shuttle between the cytoplasm and nucleus in response to changes in cellular mechanics, ECM stiffness, and cytoskeletal dynamics (Feltri et al. 2021). When the Hippo pathway is inactive, YAP/TAZ translocate to the nucleus, where they interact with transcription factors such as TEAD (TEA domain transcription factors) to promote the expression of genes involved in proliferation, differentiation, and myelination (Feltri et al. 2021; Lopez-Anido et al. 2016). Conversely, activation of the Hippo pathway phosphorylates and retains YAP/TAZ in the cytoplasm, preventing their nuclear activity. YAP/TAZ signaling is crucial for Schwann cell development, particularly in regulating their transition from immature states to fully differentiated, myelinating Schwann cells (Grove et al. 2017; Poitelon et al. 2016; Deng et al. 2017). YAP/TAZ activation has also been shown to promote Schwann cell proliferation during early development (Grove et al. 2017; Poitelon et al. 2016; Deng et al. 2017). In addition, following peripheral nerve injury, YAP/TAZ activation has been linked to Schwann cell peripheral nerve remyelination (Jeanette et al. 2021; Grove et al. 2020). Yet, while homozygous deletion of YAP in Schwann cells causes no developmental defects, deletion of TAZ causes a radial sorting delay suggesting that TAZ is more important for early Schwann cell development than YAP. In addition, TAZ can compensate for the absence of YAP (Moore et al. 2024; Poitelon et al. 2016), but YAP was not able to compensate for the absence of TAZ (Poitelon et al. 2016). However, YAP and TAZ are functionally redundant, as deletion of one YAP allele in the context of TAZ deletion aggravates the developmental defects (Poitelon et al. 2016; Grove et al. 2017). These defects are mediated by the role of YAP/TAZ-TEAD1 on Schwann cell proliferation and gene regulation by cooperating with SOX10 to activate enhancers of myelin-related genes, including EGR2/KROX20, myelin proteins, and lipid metabolism regulators (Lopez-Anido et al. 2016; Poitelon et al. 2016). Yet, YAP/TAZ do not function in isolation but instead integrate signals from multiple mechanotransduction pathways. For example, YAP/TAZ activity is influenced by Rho GTPase signaling (Weaver et al. 2025). Another important crosstalk occurs between YAP/TAZ and mechanosensitive ion channels such as Piezo channels. Piezo1-mediated calcium influx has been implicated in regulating YAP/TAZ activation, suggesting that ion channel signaling contributes to YAP/TAZ-mediated mechanotransduction in Schwann cells (Acheta, Bhatia, et al. 2022).

One emerging therapeutic approach to stimulate mechanosensitive pathways is low-intensity ultrasound (LIU) (Acheta et al. 2021). LIU, particularly in its pulsed form known as Low-Intensity Pulsed Ultrasound, is a therapeutic modality that utilizes mechanical acoustic waves at low intensities to stimulate biological tissues. Traditionally, LIU has been employed to enhance bone healing and has demonstrated efficacy in promoting regeneration and exhibiting anti-inflammatory effects across various tissues (Jiang et al. 2019). Several physical processes may stimulate the beneficial effect of LIU, including thermal effect, acoustic cavitation, and acoustic forces. First, the thermal effect of LIU is reduced when low intensity is applied in a pulsed manner, thus the thermal homeostasis in tissue limits the possibility of modulation due to temperature (Acheta et al. 2021). Second, the acoustic cavitation, to induce the formation and collapse of microbubbles in tissue, has well-supported evidence of enhancing drug delivery by increasing tissue permeability, but has not demonstrated any clear effects on control animals (Lopez-Aguirre et al. 2024). Thus, it is highly likely that the LIU effect in regeneration is mediated by acoustic forces stimulating mechanosensitive pathways in cells. A few studies have started to investigate the vibratory effect of LIU on the cell surface as a stimulus for mechanically sensitive membrane surface receptors, including extracellular matrix receptors, such as integrin (Acheta et al. 2021) or mechanosensitive ion channels (Zhang et al. 2021; Liu et al. 2022). A few studies have also shown that LIU application could modulate the expression or activation of YAP/TAZ in various cell types (smooth muscle cells, periodontal ligament cells, endothelial cells, retinal ganglion cells, and chondrocytes) (Liu et al. 2023; Jian et al. 2023; Xu et al. 2018; Zhou et al. 2018; Pan et al. 2024). In this study, we investigate whether mechanosensitive pathways in Schwann cells are activated by the application of LIU.

## 2. Methods

### 2.1 Primary Cell Culture

Sprague Dawley rats were used for the preparation of primary Schwann cell and dorsal root ganglion cultures. Primary rat Schwann cells were isolated from P3 rat pups as described in (Poitelon and Feltri 2018) and grown in high glucose DMEM supplemented with 10% fetal bovine serum, 2 mM L-glutamine, 2 μM forskolin, 2 ng/mL recombinant human neuregulin-1 (RhNRG), penicillin, and streptomycin. Primary Schwann cells in passages 3 to 7 were used. Primary dorsal root ganglion neurons were isolated from E15.5 rat embryos as described in (Catignas et al. 2021). Non-neuronal proliferative cells were eliminated by treating the culture with 10 μM Fluorodeoxyuridine and 10 μM Uridine. Under starvation conditions, Schwann cells were incubated for 12 hours in medium deprived of FBS, neuregulin, and forskolin. For verteporfin treatment (Sigma, SML0534), verteporfin was solubilized in DMSO at 2 mM, then Schwann cells were treated with either 0.5% DMSO or 10 uM verteporfin for 4 hr (unless indicated otherwise).

### 2.2 Low-intensity ultrasound treatment

For the application of LIU, Schwann cells were seeded at an initial density of 10^5^ cells in 35 mm dishes. The ultrasound transducer (a 2 cm^2^ applicator) was immersed in the media, without touching the bottom of the well. LIU parameters were set at 1 MHz, 20% pulse ratio (i.e., ON for 200 milliseconds and OFF for 800 milliseconds), and ultrasound was applied for 5 min daily (unless indicated otherwise) at the indicated intensities. Conditioned media and cells were harvested, or cells were immunostained 1 hr or 24 hrs after the last application of LIU (unless indicated otherwise).

### 2.3 Animals

This study was carried out by the principles of the Basel Declaration and recommendations of ARRIVE guidelines issued by the NC3Rs and approved by the Albany Medical College Institutional Animal Care and Use Committee (no. 20–08001). *Taz* and *Yap* floxed mice in C57BL6/129 mixed background, and P0-Cre transgenic in the congenic C57BL6/J background were described previously and labelled as *Yap*^cHet^ ; *Taz*^cKO^ (Poitelon et al. 2016). The resulting mutant mice were compared to control littermates. Males and females were included in the study. No animals were excluded from the study. Animals were housed in cages of 5 animals in 12 hrs light / dark cycles.

### 2.4. Immunohistochemistry and immunoblotting

Immunocytochemistry and immunoblotting were performed as described (Acheta, Hong, et al. 2022; Hong, Garfolo, et al. 2024). Ten micrograms of protein were used for Western blots. The antibodies used are the following: anti-YAP 1:200 (Santa Cruz, sc-101199), anti-phospho-YAP 1:200 (Cell Signaling, 4911S), anti-TAZ 1:1000 [WB] and 1:300 [IHC] (Proteintech, 23306-1-AP), anti-phospho-TAZ 1:200 (Cell Signaling, 59971S), anti-AKT 1:1000 (Cell Signaling, 9272), anti-phospho-AKT 1:1000 (Cell Signaling, 9271), anti-Pan TRK 1:8000 (Abcam, ab181560), anti-phospho-TRK (Cell Signaling, 9141), anti-calnexin 1:3000 (Sigma, C4731), anti-phospho-histone H3 1:400 (Millipore, 06-576), anti-KI67 1:1000 (Abcam, ab15580). TUNEL assays were performed on coverslips of culture as described in (Belin et al. 2019).

### 2.5. RNA Preparation and Real-Time Quantitative-PCR

Total RNA was prepared from Schwann cells with TRIzol (Roche Diagnostic). One microgram of RNA was reverse transcribed using High-Capacity cDNA Reverse Transcription Kit (Applied Biosystems, 4368814) according to manufacturer’s instructions. Quantitative PCR (qPCR) was performed similarly to (Poitelon et al. 2018) using the 20 ng of cDNA combined with SYBR Green PCR Master Mix (Applied Biosystems, 4309155). Data were analyzed using the threshold cycle (Ct) and 2(−ΔΔCt) method. RPS20 was used as an endogenous reference gene. Primers used: NGF (F, GGCCACTCTGAGGTGCATAG; R, GTCTCCCTCTGGGACATTGC), BDNF (F, AGTATTAGCGAGTGGGTC; R, GTTCCAGTGCCTTTTGTC), NTF3 (F, GGTTGCAGGGGGATTGAT; R, TATTCGTATCCAGCGCCA) and RPS20 (F, TCTGCTCTGTCCTGACTCACC; R, CTGAGGCGTAGCTTCCTGAC).

### 2.6. Statistical Analyses

Data collection and analysis were performed blindly to the conditions of the experiments. Data are presented as mean ± standard deviation (S.D.). No statistical methods were used to predetermine sample sizes, but our sample sizes are reflective of those used in existing literature within the field. Two-tailed Student’s t-test and Two-Way with Bonferroni’s post hoc test were used for statistical analysis of the differences between two or multiple groups, respectively. Values of *p* ≤ .05 were considered to represent a significant difference.

## 3. Results

### 3.1 LIU affects the adhesion and proliferation of Schwann cells

Through the review of the literature, depending on the intensity of exposure, the therapeutic ultrasound can be divided into two groups: low-intensity ultrasound (<3 W/cm^2^) and high-intensity ultrasound (≥3 W/cm^2^) (Acheta et al. 2021). Here, we used parameters for LIU intensities between 0.1 -1W/cm^2^ delivered through a pulse wave to minimize heat generation; at a 1MHz intensity to allow sound waves to penetrate 3-5 cm in biological tissues (Acheta et al. 2021). Surprisingly, we observed that when applied at 1W/cm^2^ after 5 minutes of treatment, the number of Schwann cells was significantly reduced by 75% compared to Schwann cells that did not receive the LIU treatment (Fig. 1A). We observed similar results after 10 minutes of treatment (Fig. 1B). We also show that this reduction was not associated with an increase in cell apoptosis, which suggests that application of LIU at 1W/cm^2^ may cause detachment of Schwann cells from coverslips (Fig. 1C). However, when applied at 0.1 or 0.3W/cm^2^, LIU did not decrease Schwann cell number (Fig. 1). We also controlled the effect of LIU on the temperature of the media over time and did not detect any significant change in temperature between the non-treated and LIU-treated media (Fig. 1D).

**Fig. 1.**
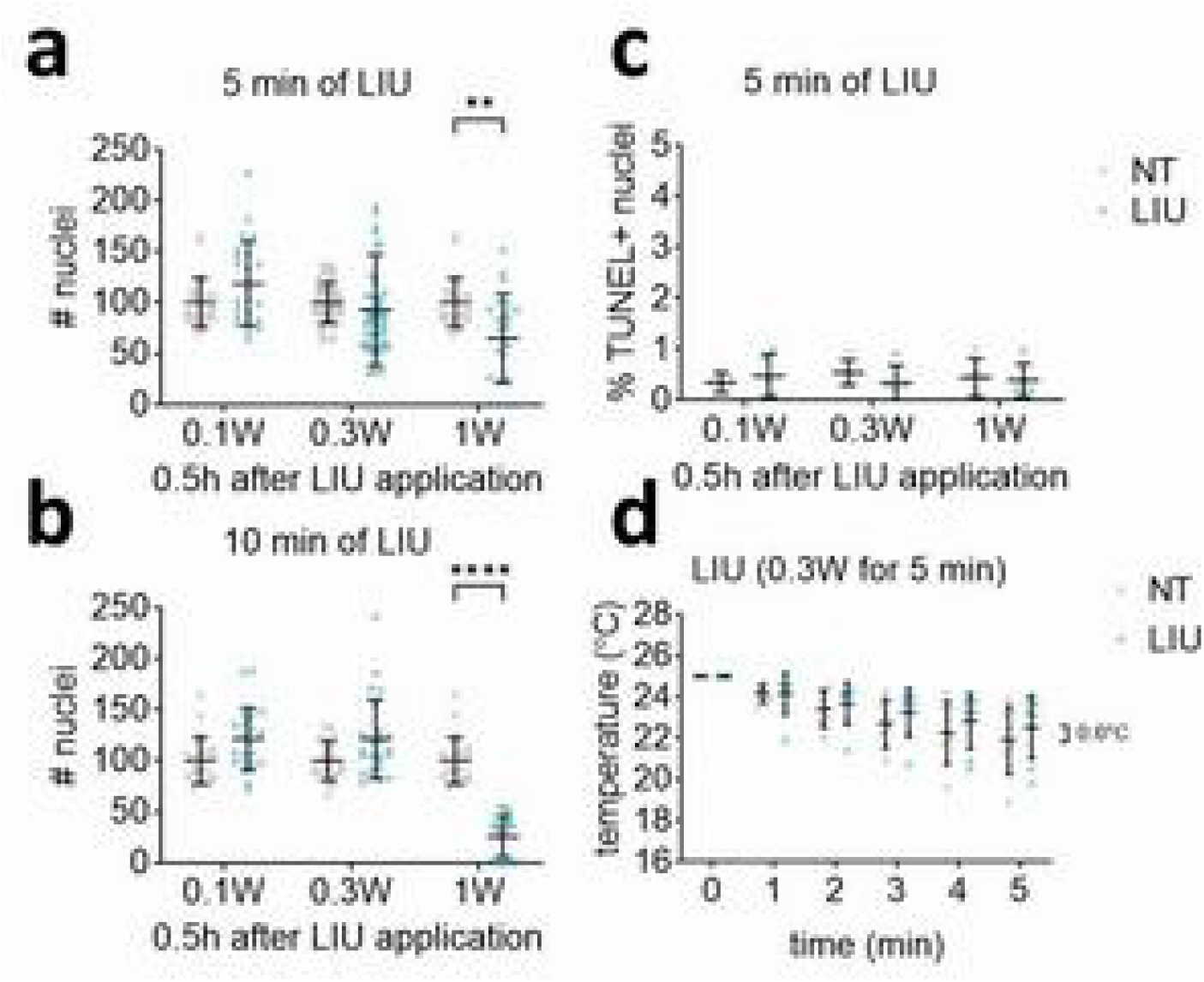
LIU application does not have detrimental effects on Schwann cell culture. (a, b) Number of Schwann cells following no LIU treatment (NT) or 5 min (a) or 10 min (b) of LIU treatment at 0.1W, 0.3W, and 1W. (c) Number of TUNEL+ Schwann cells following no LIU treatment (NT) or LIU treatment (0.1W, 0.3W, and 1W). (d) Schwann cell media temperature following no treatment (NT) or LIU treatment at 0.3W. n ≥ 3 experiments for each condition. Data are presented as means ± SD. Two-way ANOVA with Bonferroni post hoc test, ****, *p* ≤ .0001 **, *p* ≤ .01.

A recurrent observation in the literature suggests that the application of LIU increases cell proliferation (Acheta et al. 2021). Here we show that the application of LIU for 5 min at 0.3W/cm^2^ increases Schwann cell proliferation as shown by an increase in the number of KI67-positive and pH3-positive Schwann cells when compared to Schwann cells that did not receive the LIU treatment (Fig. 2A, B, E, F). Interestingly, the proliferative effect of LIU application observed at 0.3W/cm^2^ was not seen at 1W/cm^2^. This suggests either that LIU has an extremely narrow range of effective intensities or that loss of Schwann cells observed at 1W/cm^2^ (Fig. 1A) preferentially affected Schwann cells entering or undergoing mitosis, which are stages during which cells exhibit low adherence in culture (Fig. 2B, F). We also observed that the effect of LIU application on proliferation is transient, being detectable 30 min after application, but is not statistically significant at 48 hrs post LIU (Fig. 2C, G). Finally, the effect of LIU on increasing Schwann cell proliferation is obtained in the presence of growth factors, unsynchronized/unstarved conditions. However, in the absence of growth factors/starved conditions, while on average 20% of Schwann cells are still proliferating, LIU is not mediating more proliferation (Fig. 2D, H). Overall, when applied at 0.3W/cm^2^ for 5min, LIU has no apparent detrimental effect on Schwann cell primary cultures but transiently promotes their proliferation in the presence of growth factors.

**Fig. 2.**
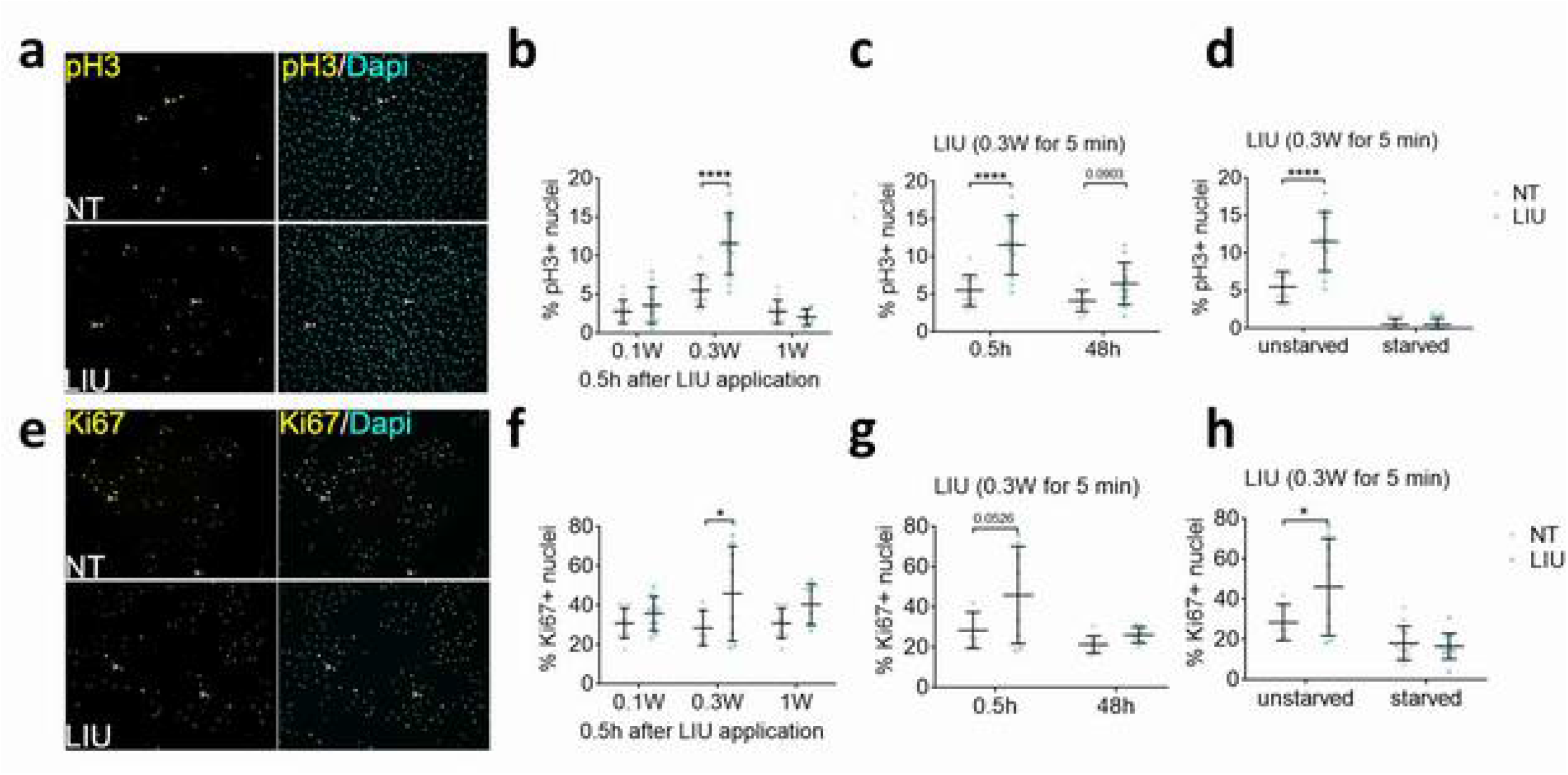
LIU application transiently increases Schwann cell proliferation. (a-h) Representative Immunostainings (a, e) and quantification (b-d, f-h) of pH3+ (a-d) and Ki67+ (e-f) Schwann cell nuclei. Quantification of Schwann cell proliferation following LIU treatment at increasing intensities (0.1W, 0.3W, 1W) and 0.5h post-application (b, f). Long-term effects on Schwann cell proliferation of 0.3 W LIU at 0.5h and 48h post-application (c, g). Effects of 12 hrs starvation and LIU treatment on Schwann cell proliferation at 0.5 h post-application (d, h). n ≥ 3 experiments for each condition. Data are presented as means ± SD. Two-way ANOVA with Bonferroni post hoc test, ****, *p* ≤ .0001; *, *p* ≤ .05.

### 3.2 LIU increases TAZ activation in Schwann cells

Activation of the Hippo pathway ultimately leads to the phosphorylation and subsequent cytosolic sequestration and/or degradation of the Hippo pathway transducers YAP and TAZ. YAP and TAZ are the main effectors of the Hippo signaling pathway and regulate the expression of a large number of genes by interacting TEA DNA-binding proteins transcription factors (Feltri et al. 2021). YAP and TAZ proliferative functions are critical to Schwann cell maturation during development and for nerve repair after injury (Grove et al. 2017; Poitelon et al. 2016; Deng et al. 2017). In addition, we have previously shown that YAP and TAZ respond to mechanical stimuli in Schwann cells and can be visualized by immunohistochemistry through the translocation of TAZ into the nucleus or by western blot through the decrease in the phosphorylated/inactive form of YAP and TAZ (Poitelon et al. 2016). Here we demonstrate that following 1 hr post application of LIU at 0.3W/cm^2^ TAZ phosphorylation decreases (as seen by an increase of the TAZ/p-TAZ ratio) and TAZ nuclear localization increases (as seen by an increase in the TAZ nuclear/cytosolic ratio) in cultured Schwann cells (Fig. 3A, B). Contrastingly, we also observe a decrease in the YAP/p-YAP ratio, suggesting that LIU modulates TAZ and YAP differently by increasing TAZ and by reducing YAP activation. Overall, these data indicate that the application of LIU has distinct effects on YAP and TAZ and suggest that LIU activates TAZ in Schwann cells.

**Fig. 3.**
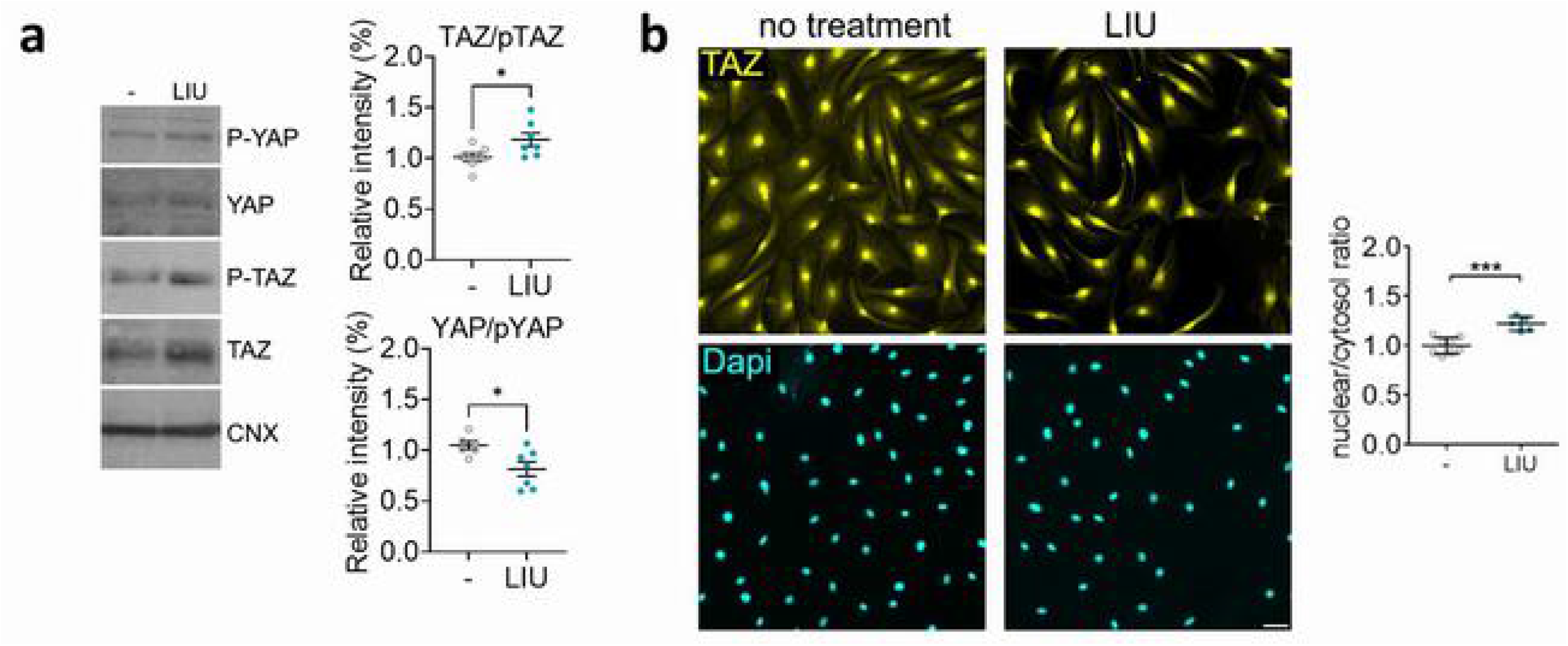
LIU application increases TAZ activation in Schwann cells. (a) Western blot analysis of phosphorylated (inactive) and non-phosphorylated (active) forms of YAP and TAZ in Schwann cell cultures following no LIU treatment (-) or LIU treatment. Data are presented as means ± SD. (b) Immunofluorescence labeling of TAZ and quantification of the nuclear/cytosolic ratio and cytosolic intensity of TAZ in Schwann cell cultures following no LIU treatment (-) or LIU treatment. Two-sided unpaired t-tests, ***, *p* ≤.001; *, *p* ≤.05.

### 3.3 LIU promotes nerve growth factor expression in Schwann cells and sensory neurons

A couple of studies have investigated the effect of LIU application on Schwann cells *in vitro* and shown an upregulation of myelination master regulator EGR2 and myelin protein gene expression (Yue et al. 2016; Huang, Jiang, et al. 2023). However, the exact molecular machinery that mediates the biomechanical effect of LIU on Schwann cells is unclear. We and others have recently shown that YAP and TAZ, together with TEAD transcription factors, are regulators of EGR2 and myelin protein expression (Lopez-Anido et al. 2016; Poitelon et al. 2016; Grove et al. 2017; Grove et al. 2024; Hong, Kirkland, et al. 2024), suggesting implications for YAP and TAZ mechanotransduction in mediating LIU effect on Schwann cells. To test if the effect of LIU application on gene expression was mediated by YAP and TAZ, we decided to focus on the expression of neurotrophins due to their critical role in the Schwann cell repair phenotype. Recent works have shown an association between the expression of neurotrophins and the Hippo pathway (Huang, Li, et al. 2023; Wang et al. 2021). In addition, several studies have suggested that neurotrophins might be able to be regulated by LIU in Schwann cells, yet results are not consistent (Ren et al. 2018; Zhang et al. 2009). First, we analyzed the expression of neurotrophic factors in mice sciatic nerves lacking TAZ and haplosufficient for YAP in Schwann cells (*Yap*^cHet^ ; *Taz*^cKO^). We used mice at 5 days of age and looked at the expression of nerve growth factor (*NGF*), brain-derived neurotrophic factor (*BDNF*), glial-derived neurotrophic factor (*GDNF*), and neurotrophin-3 (*NTF3*) in sciatic nerves. We show that the absence of TAZ in Schwann cells leads to a reduction of *NGF, BDNF*, and *GDNF* in sciatic nerves when compared to control nerves (Fig. 4A). To assess if the application of LIU was able to upregulate neurotrophins, we used in vitro primary Schwann cells and dorsal root ganglia sensory neurons cultures to independently apply LIU. We applied for LIU for 7 days and collected Schwann cells and dorsal root ganglia sensory neurons 24 hrs after the last LIU application. We demonstrate that the application of LIU leads to a 61% increase in NGF mRNA level in LIU-treated Schwann cells when compared to non-treated Schwann cells (Fig. 4B). In addition, we demonstrate that the application of LIU leads to a 45% increase in NGF mRNA levels in LIU-treated dorsal root ganglia sensory neurons when compared to non-treated neurons (Fig. 4C).

**Fig. 4.**
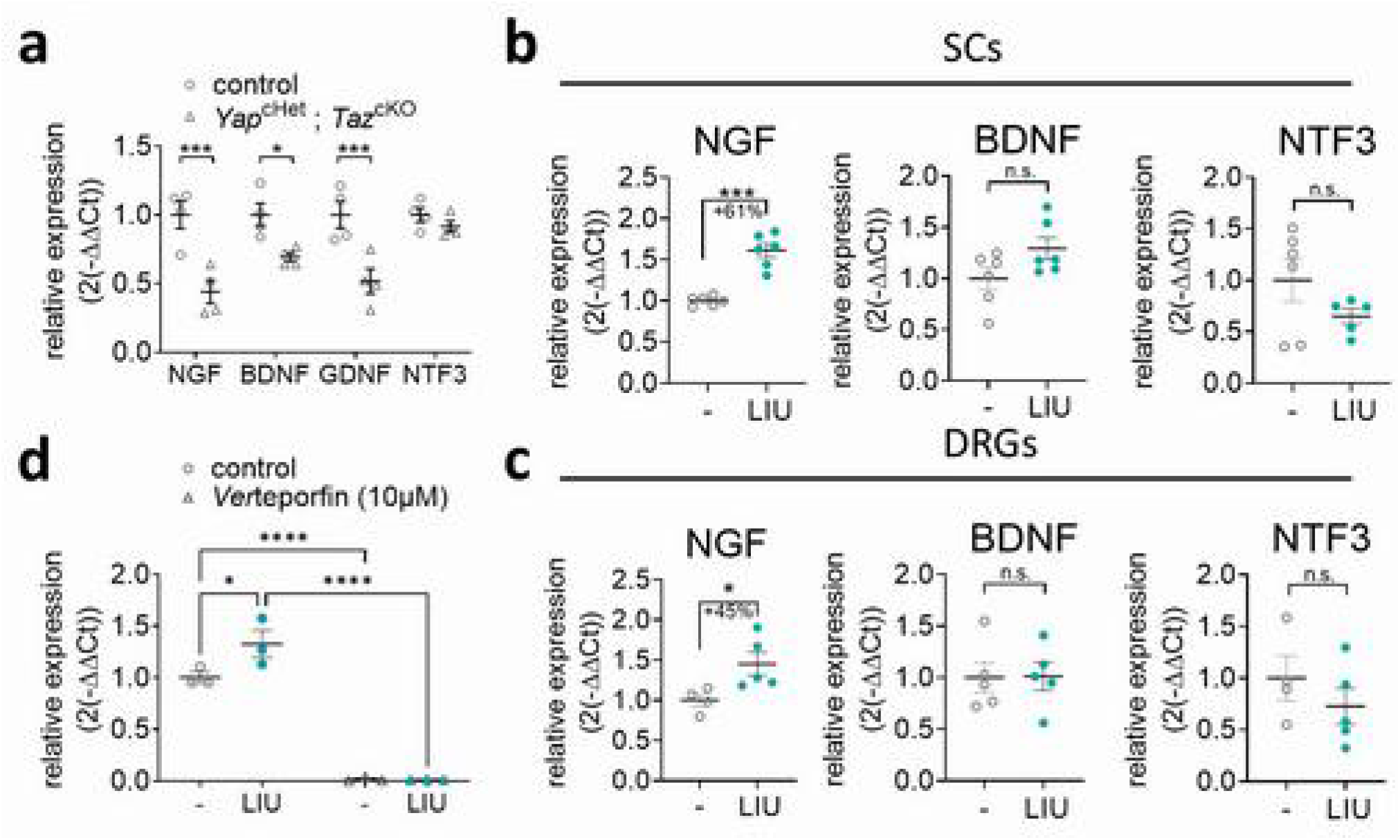
LIU application promotes nerve growth factor expression. (a) Real-time quantitative PCR of NGF, BDNF, GDNF, NTF3 in control and *Yap*^cHet^ ; *Taz*^cKO^ sciatic nerves. RSP20 was used as endogenous control. n ≥ 4 mice. Data are presented as means ± SEM. Two-way ANOVA with Bonferroni post hoc test, ***, *p* ≤ .001; *, *p* ≤ .05. (b, c) Real-time quantitative PCR of NGF, BDNF, NTF3 without treatment or with LIU treatment in Schwann cells (b) and in dorsal root ganglia sensory neurons (c). RSP20 was used as endogenous control. (d) Real-time quantitative PCR of NGF without treatment or with LIU treatment and with or without 10µM Verteporfin treatment in Schwann cells. RSP20 was used as endogenous control. n ≥ 3 experiments per condition. Data are presented as means ± SD. Two-sided unpaired t-tests (a-c) and two-way ANOVA with Bonferroni post hoc test (d), ****, *p* ≤ .0001; ***, *p* ≤.001; *, *p* ≤.05.

In our prior studies, we demonstrated that loss of YAP, TAZ, or both leads to changes in gene expression in Schwann cells through modulation of TEAD transcription factor activity (Lopez-Anido et al. 2016; Moore et al. 2024; Poitelon et al. 2016). To determine whether the YAP/TAZ-TEAD1 complex mediates the up regulation of *NGF* following LIU, we treated Schwann cells with verteporfin, a drug that disrupts YAP/TAZ-TEAD1 complex formation (Fig. 4D). Schwann cells were subjected to 5 min of LIU, followed by treatment with verteporfin or vehicle control for 4 hrs. Here, we show that verteporfin treatment abolishes NGF expression in Schwann cells. Furthermore, although *NGF* expression was increased 4 hrs post treatment with LIU, this LIU-induced upregulation was eliminated in the presence of verteporfin in Schwann cell cultures. These findings indicate that *in vitro NGF* expression in Schwann cells is regulated by YAP/TAZ-TEAD1 transcriptional complex, and that verteporfin can disrupt this regulation. Overall, our findings underscore the pivotal role of YAP/TAZ and TEAD1 in modulating *NGF* expression and highlight the therapeutic potential of targeting this pathway in LIU-based interventions.

## 4. Discussion

Our study demonstrates that LIU modulates Schwann cell behavior by influencing adhesion, proliferation, and activation of the Hippo signaling pathway. Specifically, we show that *in vitro* LIU application at 0.3W/cm^2^ transiently increases Schwann cell proliferation, whereas higher intensity (1W/cm^2^) results in cellular detachment without increasing apoptosis. Furthermore, we provide *in vitro* evidence that LIU selectively activates TAZ but not YAP, as indicated by increased TAZ nuclear localization and a decreased phosphorylated-to-total TAZ ratio. Finally, we reveal that LIU application upregulates nerve growth factor (NGF) expression in both Schwann cells and sensory neurons in culture, supporting the hypothesis that LIU promotes neurotrophic support. These findings highlight the potential of LIU as a mechanotherapeutic approach to modulate Schwann cell function and enhance peripheral nerve regeneration through neurotrophic factors.

### 4.1 TAZ is activated in Schwann cell following LIU application

One of the key contributions of our study is the mechanistic correlation between LIU stimulation and Hippo pathway regulation in Schwann cells. Our results align with prior studies that demonstrate that mechanical cues influence YAP and TAZ dynamics, and here we extend this knowledge by identifying LIU as a novel modulator of this pathway (Poitelon et al. 2016; Weaver et al. 2025).

While YAP and TAZ are often considered functionally redundant in mechanotransduction, our data suggests that LIU differentially regulates these two transcriptional coactivators, favoring TAZ activation. This is particularly intriguing given previous findings that TAZ, but not YAP, plays a critical role in early Schwann cell development, myelination, and remyelination (Jeanette et al. 2021; Poitelon et al. 2016; Moore et al. 2024). Yet because prior studies have shown TAZ activity was able to compensate for YAP loss of function, it is unclear if the activation of TAZ we measured is a direct response to LIU stimulation or a compensatory molecular response following LIU-mediated YAP reduced activation. Further investigation into the molecular mechanisms governing YAP and TAZ specific response to LIU could provide valuable insights into Schwann cell mechanobiology.

### 4.2 Application of LIU promotes SC proliferation and production of nerve growth factors

Our study provides new insights into the role of mechanotransduction in neurotrophic factor expression. Previous *in vitro* studies have reported inconsistent effects of LIU on neurotrophin levels in Schwann cells, with some suggesting increased expression and others reporting no significant changes (Ren et al. 2018; Zhang et al. 2009; Li et al. 2023). Neurotrophic factors have been shown to promote neuroprotection, axonal regrowth, and even myelinogenesis following peripheral nerve injury (Boyd and Gordon 2003), with NGF being essential for sensory neurons (Barker et al. 2020) while BDNF and NTF3 are predominantly involved with motor neuron survival and function (Stansberry and Pierchala 2023). Here, we corroborate prior evidence showing that specific neurotrophic factors (*i*.*e*., NGF) are increased in cultured Schwann cells following LIU application. Importantly, we found that the effect of LIU on NGF expression is not limited to Schwann cells but is also present when LIU was applied to sensory neurons, thus suggesting that the effects of LIU on NGF expression are mediated by a common mechanosensitive pathway present in both cell types. Several signaling pathways are known to regulate NGF, namely CREB signaling which increases NGF expression in response to neuronal activity, c-Fos/c-Jun signaling which increases NGF expression in response to injury and inflammation (Mullenbrock, Shah, and Cooper 2011). Specifically, we demonstrated that treatment with verteporfin, an inhibitor of YAP/TAZ-TEAD1 interaction, abolishes NGF expression in Schwann cells, even in the presence of LIU. This indicates that the YAP/TAZ-TEAD1 complex is essential for *NGF* upregulation in response to mechanical stimulation with LIU. Thus, further research is necessary to determine whether TEAD’s role in NGF transcriptional regulation is direct, through TEAD binding elements in NGF, or indirectly through the regulation of *NGF* regulators. Given the essential role of NGF in neuronal survival and axonal regeneration, our findings suggest that LIU-based interventions hold potential for enhancing the repair of injured peripheral nerves.

### 4.3 Concluding Remarks

Our study advances the understanding of mechanotransduction in Schwann cells by identifying LIU as a novel stimulus for TAZ activation, proliferation, and neurotrophin expression. These findings provide a foundation for future research exploring LIU as a potential therapeutic tool for peripheral nerve regeneration. While our results establish a mechanistic framework linking LIU to Schwann cell function, additional studies are needed to determine the long-term effects of LIU treatment in vivo and to optimize stimulation parameters. Future research should also investigate the interplay between LIU and other mechanotransduction pathways, including integrin signaling and mechanosensitive ion channels, to further refine our understanding of the molecular mechanisms underlying LIU-induced Schwann cell responses.

## Acknowledgments

This work was funded through grants from the National Institute of Neurological Disorders and Stroke (R01NS110627) awarded to Y.P. and (R01NS134493) awarded to S.B. J.A. was supported by a NIH Diversity supplement on R01NS110627.

